# Intestinal interstitial fluid isolation provides novel insight into the human host-microbiome interface

**DOI:** 10.1101/2024.01.11.574524

**Authors:** Ellen G. Avery, Lea-Maxie Haag, Victoria McParland, Sarah M. Kedziora, Gabriel J. Zigra, Daniela S. Valdes, Marieluise Kirchner, Oliver Popp, Sabrina Geisberger, Olivia Nonn, Tine V. Karlsen, Gabriele N’Diaye, Alex Yarritu, Hendrik Bartolomaeus, Theda U.P. Bartolomaeus, Moritz I. Wimmer, Nadine Haase, Andreas Wilhelm, Gerald Grütz, Olav Tenstad, Nicola Wilck, Sofia K. Forslund, Robert Klopfleisch, Anja A. Kühl, Raja Atreya, Stefan Kempa, Philipp Mertins, Britta Siegmund, TRR241 IBDome Consortium, Helge Wiig, Dominik N. Müller

## Abstract

**Aims:** The gastrointestinal (GI) tract is composed of distinct subregions which exhibit segment-specific differences in microbial colonization and (patho)physiological characteristics. Gut microbes can be collectively considered as an active endocrine organ. Microbes produce metabolites, which can be taken up by the host and can actively communicate with the immune cells in the gut lamina propria with consequences for cardiovascular health. Variation in bacterial load and composition along the GI tract may influence the mucosal microenvironment and thus be reflected its interstitial fluid (IF). Characterization of the segment-specific microenvironment is challenging and largely unexplored because of lack of available tools.

**Method and Results:** Here, we developed methods, namely tissue centrifugation and elution, to collect IF from the mucosa of different intestinal segments. These methods were first validated in rats and mice, and the tissue elution method was subsequently translated for use in humans. These new methods allowed us to quantify microbiota-derived metabolites, mucosa-derived cytokines, and proteins at their site-of-action. Quantification of short-chain fatty acids showed enrichment in the colonic IF. Metabolite and cytokine analyses revealed differential abundances within segments, often significantly increased compared to plasma, and proteomics revealed that proteins annotated to the extracellular phase were site-specifically identifiable in IF and were differentially expressed when compared to matched serum, all suggesting local synthesis.

**Conclusion:** Collection of IF from defined segments and the direct measurement of mediators at the site-of-action in rodents and humans bypasses the limitations of indirect analysis of fecal samples or serum, providing direct insight into this understudied compartment.

## Introduction

The gut microbiota consists of a complex community of bacteria that is essential for immune homeostasis and has important implications for host health.^1^ It is considered as an endocrine organ, producing metabolites that interface with host physiology by triggering responses in the local intestinal microenvironment or in distant target organs.^2^ Microbiomes from diseased and non-diseased individuals differ (exhibiting a *dysbiotic* as opposed to *eubiotic* state), as shown for several inflammatory conditions (e.g. colitis), hypertensive CVD, and metabolic disorders.^3–13^ Low-grade inflammation can be triggered through dysbiosis and its derived metabolites.^14^ Although it is well recognized that there is considerable variability in the microbes that colonize the intestinal segments, current microbiome research mainly focuses on fecal samples, thereby losing segment-specific information. The use of feces to study the segment-specific host-microbiota interactions may therefore be considered as disadvantage, as this sample represents only the total end-product of a highly dynamic system. Thus, we aimed to develop methods to collect interstitial fluid (IF) from different segments of the intestinal mucosa of rodents and humans to gain direct insight into the site-specific microenvironment of the gut.

IF is simply defined as the fluid found in the spaces between the cells of a tissue, the tissue’s structural components known as the extracellular matrix (ECM), and the capillaries.^15^ The extracellular fluid space consists primarily of IF and a small percentage of plasma.^16^ Isolated IF has been used to reveal tissue-specific microenvironmental signatures present in the extracellular space which cannot be predicted by serum measurements.^17^ demonstrating the importance of measuring IF content locally at the site of inflammation.

The GI tract is the interface between the host and its microbiota. We assume that the intestinal IF is rich in both host-produced and microbially produced factors such as cytokines, metabolites, and proteins, although this has not yet been demonstrated. Little is currently known about the spatial composition in the intestinal mucosal interstitium, the interaction with immune cells and the consequences for health and disease. To overcome the limitations of existing approaches, we aimed to develop and validate new methods that can be applied to rodent and human biopsies, allowing direct phenotyping of the local intestinal microenvironment instead of indirect phenotyping of fecal samples.

## Materials and Methods

Detailed description of all analytical methods and data analysis used are available in the Supplementary material online.

## Results

### Intestinal IF can be isolated from rats by tissue centrifugation

The centrifugation technique to isolate native IF was first described for skin and tumors in rats.^18^ To investigate whether this method could be applied to extract IF from the gut interstitium in rats, we excised the entire gut immediately after euthanasia. Gut segments were placed on a nylon mesh and subjected to 400 *g* for 10 min (Figure 1A). Following centrifugation, the volume was determined, and the isolated fluid was aliquoted for further analysis. The average fluid yield from the different segments ranged from 5 μl in duodenum to 25 μl in the other segments, respectively (Figure 1 B, C).

**Figure 1.**
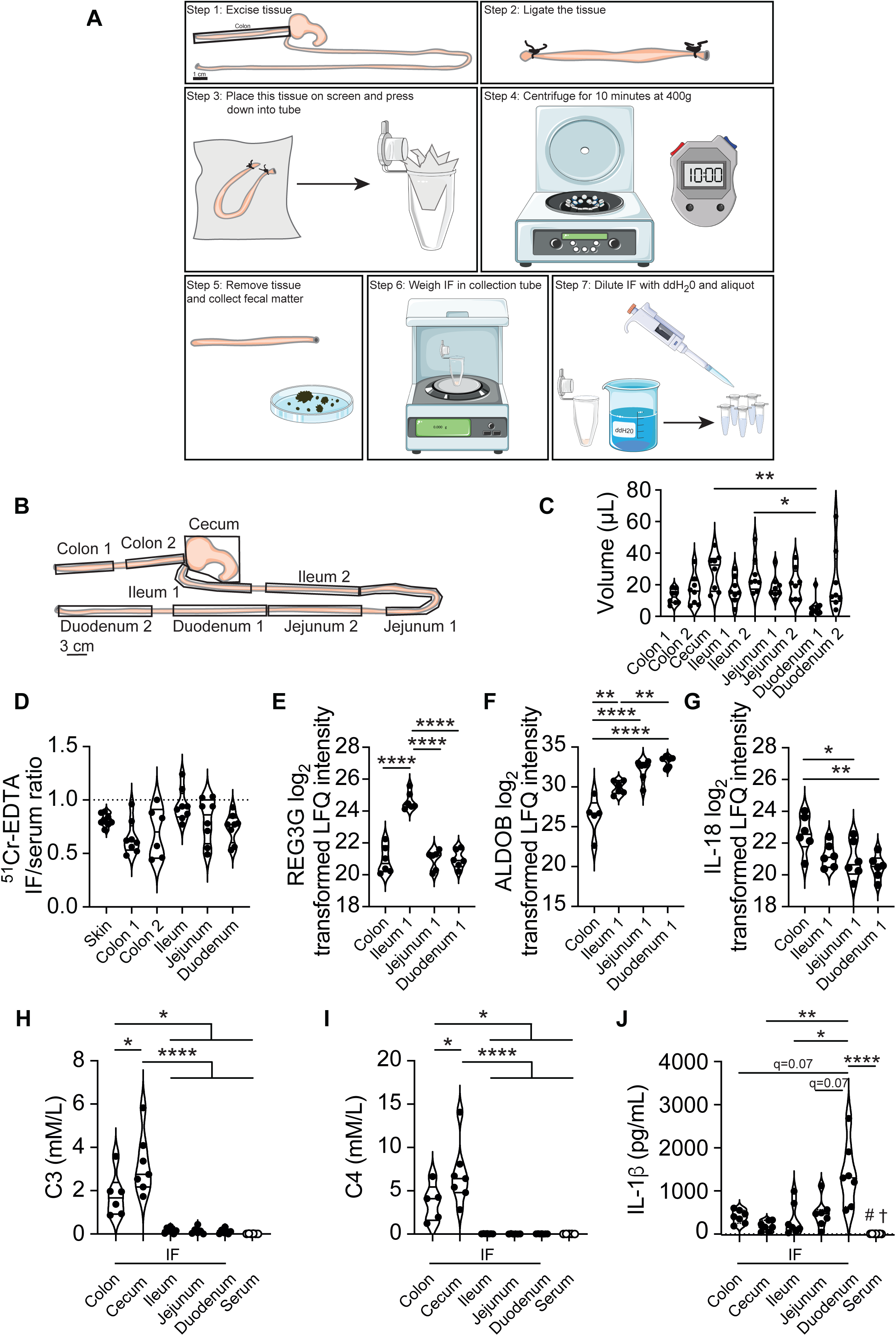
Centrifugation-based native extracellular fluid isolation within the GI tract of Sprague-Dawley rats. A) Schematic of the centrifugation-based method for IF isolation. The GI tract was excised from just below the stomach to the colon. Seven to10 cm pieces of GI tissue were measured, cut and immediately ligated. The tissue was inspected for holes, and any fecal matter was squeezed out from the edges of the tissue after the ligation point. Tissue ends were trimmed if necessary. The tissue was then placed on a light screen material, folded, and in a 5 mL Eppendorf tube. Closed tubes were centrifuged for 10 minutes at 400 x g. Immediately after removal of the tissue, the ligations were cut off and the fecal matter was collected. The IF from within the 5 mL Eppendorf tube was weighed, and diluent was added for aliquoting the centrifuged IF. IF was stored at −80 °C for further analysis. Some images were adapted from the Servier Medical Art collection (https://smart.servier.com). B) The scheme for collection of rat sample is shown. C) Centrifugation volumes of IF from male and female rats (n=8) as collected in B). D) The extracellular tracer ^51^Cr-EDTA was used to determine how well the centrifugation method works to isolate IF from GI tissue compared to skin. The IF/serum ratio of ^51^Cr-EDTA was determined using a gamma-counter from centrifuged IF (n=8). The log_2_ difference of REG3G (E), ALDO B (F) and IL-18 (G) quantified by mass spectrometry is shown for the respective intestinal segments (n=6). REG3G is selectively enhanced in the ileum, whereas IL-18 is increased in the colon. H-J) Propionate (C3), butyrate (C4) and IL-1β levels of IF from different segments of the rat GI tract (n=5-7 per segment). Statistical analysis for D) was performed using an ordinary one-way ANOVA with Dunnett’s post-hoc test (recommended for comparison to a reference condition) applied to compare the GI tissue segment IF/serum ratio to the skin IF reference method. None of these post hoc comparisons were significant. Data were examined for outliers using the ROUT outlier test. Outliers were removed when appropriate. Data in C-I) were tested by ordinary one-way ANOVA with Tukey’s multiple comparison test; *p≤0.05, **p≤0.01, ****p<0.0001, J) with Kruskal-Wallis test corrected for multiple comparisons by controlling FDR (Benjamini, Krieger, Yekutieli) #p<0.002 vs. colon and jejunum, †p<0.05 *vs.* ileum.

The origin of the isolated fluid, i.e., whether it is derived solely from the extracellular space or is “contaminated” with intracellular fluid from tissues and blood cells, can be assessed by determining the ratio of a tracer that equilibrates only in the extracellular fluid phase and does not pass intracellularly.^18^ An example of such a tracer is the non-native labeled extracellular tracer ^51^Cr-EDTA. After a two-hour *in vivo* equilibration period of ^51^Cr-EDTA, we harvested and centrifuged the corresponding intestinal segments (Figure 1D) and calculated the tracer concentration ratio in IF (centrifuged) and plasma. In these experiments, skin was included as a reference. As shown in Figure 1D, except for the ileum, the mean of the IF/serum ratio was <1.0, indicating that there was a slight dilution of the IF with intracellular fluid. In an attempt to validate that the fluid isolated by centrifugation reflected the intestinal microenvironment, we determined, as a proof-of-concept, levels of the antimicrobial peptide regenerating islet-derived protein 3 gamma (REG3G), which is produced by Paneth cells in the ileum through interactions directed by symbiotic bacteria.^19, 20^ As expected, REG3G enrichment was specific for the ileum (Figure 1E).

The enzyme aldolase B (ALDOB) is enriched in the small intestinal segments compared to the colon (Figure 1F). Next, we focused on interleukin (IL)-18, which is expressed by enteric neurons and controls goblet cell expression of antimicrobial proteins.^21^ The frequency of goblet cells increases from the upper to the lower GI tract.^22^ IL-18 was significantly increased in the colonic IF compared to the small intestinal IF (Figure 1G). We next aimed to validate the site-specific production of short-chain fatty acids (SCFA, propionate C3 and butyrate C4) as a use case for the measurement of microbial-derived metabolites. As expected, C3 and C4 concentrations measured directly in centrifuged IF were significantly higher in the colon with a higher level in the cecum compared to the small intestine and serum (Figure 1H-I). We further asked whether the cytokine IL-1β was also produced locally in the rat intestine. Duodenal IF showed significantly higher IL-1β concentrations compared to other segments of the small and large intestine as well as serum suggesting a local production (Figure 1J). Taken together, these data indicate that native IF isolated by the centrifugation-based method in a rat model is suitable for analytical phenotyping of microbial or intestinal-derived metabolites, cytokines, or proteins.

### Isolation of intestinal IF from mice using tissue elution

Since the mouse is a preferred model in many biomedical contexts, and we therefore investigated the possibility of isolating murine IF. First, we assessed the extracellular volume (ECV) in different GI segments, as this provides information on the feasibility (Figure S1A) of extracellular tracer experiments with ^51^Cr-EDTA. The ECV of GI tissue was relatively similar to that of muscle tissue^15^, with an estimated volume of 0.1 to 0.2 mL/g wet weight, depending on the segment of interest (Figure S1B). Total tissue water (the ECV and intracellular volume [ICV]) within a given tissue was also determined from the colon, cecum, ileum, jejunum, and duodenum (Figure S1C). There were some very small but significant differences in the total tissue water for individual segments, although these are unlikely to be physiologically relevant. The maximum difference between any segments was 0.02 mL/g and the mean total tissue water value across all samples was 0.79 mL/g (Figure S1C).

These distribution volume experiments indicated that the interstitial fluid space was of a size making IF isolation by centrifugation feasible. We therefore applied the same centrifugation approach to mouse intestinal tissue. As might be expected due to the lower total tissue weight of the GI tract, IF sample volumes were smaller than in rats. Although these experiments suggested that the centrifugation method was feasible for isolation of mouse intestinal IF, the sample volume was insufficient to overcome the limit of detection for many analyses unless pooled. We therefore developed an elution-based method to phenotype the mouse compartment. Elution-based methods have previously been used to isolate IF from various tumor tissues and are advantageous because they result in a large volume of fluid which can be used for multiple measurements.^15^ The general principle of the method is that the cleaned excised tissue is placed in an isosmotic buffer to maintain tissue integrity and osmotic pressure. Substances dissolved in the IF phase will equilibrate with the surrounding buffer by diffusion over time. The eluate is diluted relative to the original IF in the native tissue, but the dilution factor is known and can be used to calculate the tissue concentration of the analytes. Schematics of the methodological details is shown in Figure 2A.

**Figure 2.**
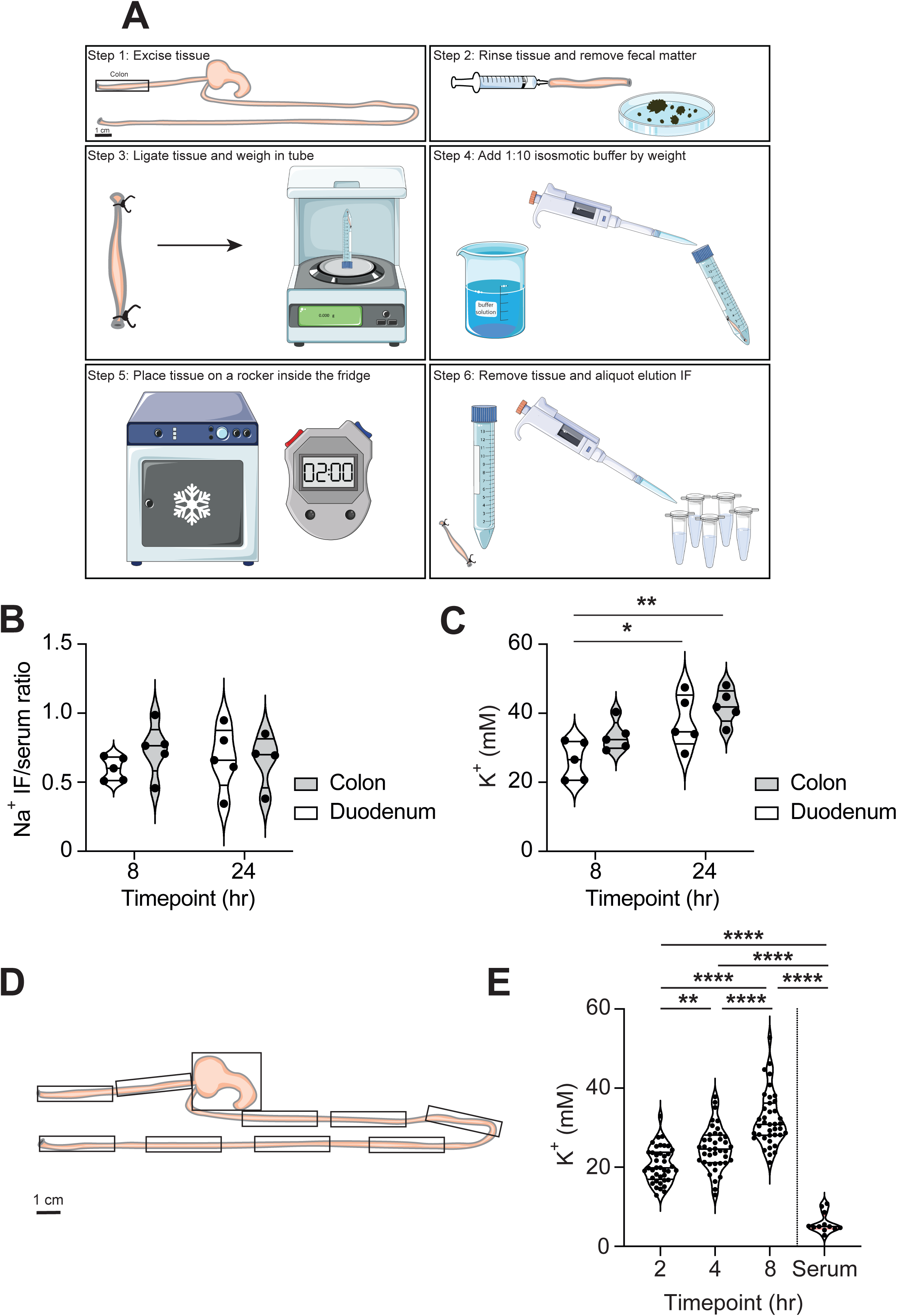
Schematic for the collection of IF using an elution-based method. A) The respective GI tract tissue was excised and immediately washed from the luminal side using a blunt syringe with approximately 5 ml of isotonic buffer to ensure the removal of any luminal contents. Fecal matter was collected from the indicated segment prior to rinsing. Both ends of the tissue were ligated to ensure that if any microscopic luminal contents were still attached to the inner surface of the gut, they would not be extracted into the surrounding buffer. After ligation, the tissue was placed in a 1:10 dilution of an appropriate isosmotic buffer solution and placed on a rocker at 4 °C. After the specified elution time, the tissue was removed, and the eluted fluid was aliquoted and stored at −80 °C for further analysis. Some images were drawn with Servier Medical Art from Servier, licensed under a Creative Commons Attribution 3.0 Unported License (https://smart.servier.com). B) Segments were taken from each region of the colon and duodenum and were either placed in mannitol for 8 or 24 hours, respectively. Comparison of the Na^+^ IF to serum ratio (C) and IF K^+^ (D) by ion chromatography from samples eluted in a mannitol-based buffer for 8 or 24 hours. N=5 for all conditions, and one outlier was excluded. D) The segmentation approach from C57BL/6J mice. E) K^+^ was measured from IF and serum (matched by animal). Separate animals were processed for each time point, n=4 for each of the 10 segments at each timepoint. The results from each time point were binned, and the results from each segment are shown together. Shorter time points result in significantly less K^+^ (mM) in elution IF. Two-way ANOVA and post-hoc Sidak multiple comparison test were used to test significance. P values are as follows; *p≤0.05, **p≤0.01.

Saline is a typical state-of-the-art isosmotic buffer option for elution. In the initial phases of method development, the ionic composition of IF was used as a proxy to measure the efficacy of the elution method, making saline inappropriate, and we used mannitol as the eluent. Theoretically, the ionic composition of IF should closely resemble that of plasma with low levels of potassium (K^+^; 3.5-5.0 mmol/L) and levels of sodium (Na^+^) ions around 140 mmol/L^15, 23^ and thus a ratio of IF Na^+^ to serum Na^+^ close to 1.0. To optimize the use of elution from GI tissues, we tested different elution time points (8 and 24 hours) and analyzed their effect on IF ion levels (Figure 2B-C). The longer 24 h time point resulted in higher K^+^ in eluted samples ranging from 20 to 40 mmol/L (Figure 2C). Increased K^+^ levels indicate leakage of intracellular K^+^ into the eluate and/or decreased Na^+^/K^+^ ATPase activity, which may be a result of energy deficit of the eluted tissue and loss of tissue integrity. Thus, we used shorter elution time points for comparison (2, 4, and 8 hours) and repeated the experiment with a larger number of intestinal segments. Across all segments, shorter elution times resulted in less K^+^ deposited in the elution buffer (Figure 2D-E). The K^+^ level within any given elution IF sample was still far higher than serum (Figure 2E), although this was not entirely unexpected, given the strong gradient of intracellular K^+^ and high ICV relative to ECV in the GI tract. These experiments suggest that there is a significant leakage of K^+^ ions originating from the intracellular space. The IF/serum Na^+^ ratio was significantly different from 1.0 (Figure 2B), indicating that this fluid shift was accompanied by cell water.

Since ion chromatography measurements of shorter elution times suggested an improved tissue integrity, we next tested whether the substances of interest would be efficiently extracted from the intestinal IF at these lower time points. The extracellular tracer ^51^Cr-EDTA (molecular weight=339) mimics the size of a small molecule/metabolite and was used to assess the time likely required for a compound to equilibrate with the elution buffer (Figure S2). The amount of tracer in the elution buffer was assessed relative to the total amount of tracer in each respective tissue segment. ^51^Cr-EDTA equilibrated with the surrounding elution buffer rather quickly, with most of the tracer being eluted after 2 hours (Figure S2). In the small intestinal segments, there was more variation in the eluted tracer fraction. We suspect that this is due to the lower ECV in the small intestine (Figure S1B), as it is known that IF is more difficult to be mobilized from spaces with a lower IF fraction.^15^ Even at the longest time point studied, a small amount of the ^51^Cr-EDTA was not recoverable, as indicated by an eluted fraction unequal to 1.0. This suggests that there may be some unspecific binding of the ^51^Cr-EDTA within the tissue, rendering a fraction of this molecule inaccessible. Nevertheless, based on these results it is likely that small molecules such as microbiota-derived metabolites can rapidly equilibrate within the elution fluid.

### IF collected using elution can be used to study tissue-specific proteomic signature

Next, we examined whether the IF from the GI tract has a protein signature that is distinct from the circulating blood as previously shown for microenvironments such as tumors^23, 24^ and the skin.^17^ Shotgun proteomics was used to identify similarities and differences between the IF and serum from C57BL/6 mice. Overall, significantly more unique protein IDs were found in IF compared to serum (Figure 5A-B). Of these, 1802 proteins were statistically significantly different between elution IF and serum (Figure 5C). While many proteins from the serum and IF spaces overlapped (Figure 5C), 2581 were found to be unique to the IF space, of which 1404 were shared between the colon and duodenum segments. There was a much greater overlap in the identifiable proteins between these two IF groups than with the serum (Figure 5C). The normalization for these data was based on the amount of protein in the starting material rather than per volume of IF or serum. Because the protein concentration in IF is typically lower than in serum^15^ and because we measured an equal amount of several peptides from each matrix in shotgun proteomics experiments, we expect that there may be a slight overrepresentation of some proteins in the elution IF. Nevertheless, proteins which have been documented to be highly abundant in serum, such as albumin (ALB)^15^, did indeed appear to be significantly higher in serum compared to elution IF (Figure 5B). Notwithstanding this fact, IF may be a better way to study proteins relevant to GI health and disease which are present or undetectable in serum due to the increased coverage of proteins from the IF space compared to the serum. Encouragingly, within the group of proteins which were robustly identifiable from the IF, there were some candidates such as galectin-3 (LGALS3; Figure 5D)^25^ that have a known biological function in the GI tract. Antimicrobial peptides such as α-defensins (DEFA) are produced by Paneth cells. As expected, the DEFA cluster 6/9/11/15/24 was significantly enriched in mouse duodenal IF (Figure 5E). In line with this, ALDOB was also significantly enriched in the duodenum (Figure 5F) as in the native rat IF (Figure 1F). Next, we demonstrated that the elution method can detect SCFA (C3 and C4) in colonic IF (Figure 5G-H). Of note, colonic and cecal C3 levels were 150- and 280-fold higher, respectively, compared to plasma. For C4, the respective fold change was even more pronounced (200 and 900-fold). These concentrations are based on the assumption that the substances are distributed throughout the entire fluid phase (total tissue water) of 0.79 ml/g tissue. However, if they distribute only or partially in the interstitial fluid phase of 0.1-0.2 mg/g tissue, which is a reasonable assumption, their concentration will be several times higher. The same reasoning applies to eluted human samples (see below).

Next, we sought to confirm that the elution-based method recapitulated the signature observed with the centrifugation method. For these experiments, we used the SD rat model, as the larger GI tissue was suitable for direct comparisons between the two methods. Indeed, REG3G and ALDOB obtained by the elution method (Figure 4A-B) showed a similar pattern in colonic IF as in the native IF of the centrifugation method (Figure 1E-F), with significantly lower abundance compared to small intestinal segments of rats. Similarly, IF concentration of IL-18 was significantly higher in the colon using both the centrifugation and elution methods (Figure 4D, Figure 1G). Additionally, LGALS3 and ALDOB were regulated in the rat elution IF (Figure 4B-C) similar to what was shown in mice (Figure 5D and 5F, respectively). Mass spectrometry analyses of rat IF isolated by the elution method also confirmed the pattern of SCFA (propionate C3 and butyrate C4; Figure 4E-F) observed in the native IF collected by the centrifugation method.

SCFA are known to infleunce immune homeostasis. Given the high sensitivity of the proteomics method used, head-to-head comparison of the two methods initially revealed apparent method-specific clustering (data not shown). After independent z-scoring within the centrifugation and elution methods, a selection of inflammatory response and immune response annotations (a total of 285 proteins) from the rat Uniprot database is shown by PCA (Figure 4G) and Pearson’s correlation (Figure 4H). The overall proteomic signature showed a similar congruence between segments (data not shown). Segments from the small intestine overlapped to some extent, whereas the colon clustered independently from the rest of the GI tract. The correlation between segments across methods was much higher than between all segments within a given method (Figure 4G). Of note, the segment with the lowest correlation between the two methods was the ileum (Figure 4H). While the ileum correlated with the duodenum in the centrifugation method, the ileal segment correlated better with the jejunum in the elution method the ileal segment (Figure 4H). This may be due to slight differences in the excised segment, as small intestinal subregions are difficult to grossly differentiate without histologic analysis.

### Tissue elution for interstitial fluid isolation can be translated for use in humans

Having established the isolation of IF in rodents, we turned our attention to translating the procedure to human samples. Phenotyping of the human intestinal interstitium is of interest in diseases such as inflammatory bowel disease (IBD), as this is the central compartment of disease activity. To the best of our knowledge, IF phenotyping has not been done performed in the context of IBD. Ideally, a method for isolation and analysis of human intestinal IF would be implementable into clinical routine and, at best, into treatment decisions. We therefore aimed to isolate IF from human intestinal mucosa biopsies obtained during colonoscopy. Of note, a colonoscopy is preceded by a routine, standardised bowel cleansing. Routine clinical human mucosal biopsies weigh an average of 9 mg, which is < 10% of the weight of rodent biopsies. Because of these very small weights, we decided to use the elution method. In this pilot study, we enrolled two patients with ulcerative colitis (UC) and one patient with Crohn’s disease (CD). Furthermore, four patients undergoing surveillance colonoscopy served as healthy controls (HD). As shown in Figure 5A, we obtained mucosal biopsies from the highly inflamed descending colon and the terminal ileum of two patients with ulcerative colitis (UC). The patient with CD presented with a severe segmental inflammation of the colon. During colonoscopy of this patient, we were able to obtain biopsies from the inflamed colon, but not from the ileum. In addition to obtaining intestinal biopsies, we drew blood to obtain paired serum samples.

First, we performed label-free proteomics and identified protein IDs ranging from 5697 to 5897 in human colon IF samples and from 5414 to 5696 in human ileum IF samples. When we compared the proteome of colonic IF samples to that of ileal IF samples from healthy donors, we found a clear separation between the two anatomical sites (Figure 5B). Colonic IF samples showed a high similarity between healthy donors, whereas ileal IF samples showed a higher variance (Figure 5B). A total of 5967 unique protein IDs were detected in all elution IF samples from healthy human donors (Figure 5C). Among these, 319 protein IDs were significantly different between the two studied intestinal sites, including several proteins known to be relevant for human GI tract health such as REG3A, DEFA5, DEFA6 and ALDOB (Figure 5C). As expected from the anatomical origin, antimicrobial proteins were significantly more abundant in ileal than in colonic IF (Figure 5D-G).

In humans, as in rodents, the terminal ileum also contains SCFA-producing microbes.^26^ Therefore, it was not surprising that we detected propionate (C3) in the IF of both anatomical sites. Propionate levels in IF were more than 20-fold higher than in plasma (Figure 5H). Despite the cleansing procedure preceding the colonoscopy, the detected plasma levels for both propionate and butyrate (data not shown) were in a similar range compared to previous studies.^27^ However, we were only able to detect propionate, but not butyrate in our eluted fluids (data not shown). Butyrate is also metabolized by colonocytes, which may be one reason for the lack of detection on addition to the effect of the bowel cleansing prior to endoscopic examination.

Shotgun proteomics showed increased levels of S100A8 and S100A9 in IF from IBD patients, which as a heterodimer represent calprotectin (Figure 5I). Serum amyloid A (SAA1), a robust biomarker of colorectal inflammation in UC patients, was also elevated in IF from IBD patients compared to healthy donors (Figure 5I). IBD patients had increased levels of defensin alpha 5 and 6 (DEFA5 and 6), microbicidal and cytotoxic peptides involved in the defense against bacteria and viruses, which are also prognostic markers for colorectal cancer (Figure 5I). Moreover, ALDOB, a protein known to enhance fructose metabolism and provide fuel during tumor cell proliferation was increased in colonic IF from IBD patients but showed no difference compared with HD in ileal IF (Figure 5I). Of note, despite having the same underlying active disease, each of the two UC patients had a distinct signature in their colonic IF (Figure 5I). The colonic IF signature for each patient was also distinct from the corresponding paired plasma sample (Figure 5J). This demonstrates that analysis of peripheral blood samples does not reflect the inflammatory microenvironmental signature present at the site of action. While UC1 had increased IL-1α and β, IL-2, IL-4, IL-5, IL-6, IL-8, and TNFα and β, GM-CSF in the colon, the second UC patient (UC2) had increased IL-17 levels (Figure 5J). Furthermore, our CD case showed a unique TH1-like profile which was different from the UC patients, as well as a slight induction of IL-2 and IL-4 in the colonic IF (Figure 5J). Anti-TNFα therapy is one treatment option in both UC and CD. Plasma TNFα levels in UC1 and UC2 were 2.8 and 2.1 pg/mL, respectively. However, in colonic IF, TNFα levels were increased 456-fold in UC1 and 40-fold in UC2 compared to plasma. While UC1 also showed 23-fold elevated ileal TNFα levels, UC2 was only 2-fold increased compared to plasma (data not shown). It is tempting to speculate whether the differences in TNFα levels within the tissue could help predict the success of anti-TNFα therapy in these patients. Taken together, spatial analysis of intestinal IF may hold promise regarding selection and prediction of therapy.

## Discussion

The body’s mucosal barriers, which separate the external environment from the internal tissues, play a critical role in maintaining host health. These selectively permeable barriers allow the entry of certain nutrients, solutes and metabolites which can interact and influence host (patho)physiology. However, accessing the tissue microenvironment within the GI tract is challenging. Interstitial fluid representing the microenvironment within the tissue, can provide unique information about cellular processes that cannot be obtained from other fluid sources, particularly plasma. Because such fluid may not be readily available, several approaches have been applied to access the tissue microenvironment.^15^ Here, we have introduced and validated two approaches to access the interstitial fluid within the GI tract: tissue centrifugation and elution. We show that each of these methods can be used to understand cellular processes occurring in different segments of the intestinal mucosa and have succeeded in translating the IF collection method from rodents to humans. As use case, we compared IBD patients during an acute flare with healthy controls to provide proof-of-principle data on the utility of our approach. Of note, we were able to show a personalized cytokine signature in colonic IF for all of our IBD patients which we would have missed using plasma phenotyping. Taken together, these methods of IF collection have important implications for the study of the GI microenvironment and may be useful in identifying novel modulators of the microbiome-immune interface.

Sampling of IF within the tissue can be challenging regardless of the site-of-interest. When sampling is possible, prenodal lymph is likely to be the best representative of native interstitial fluid, allowing assessment of the absolute concentration, which is important in determining the origin of the fluid ^15^ (see below). Such fluid has recently been collected from specific anatomical sites in mice and rats^28^. The sampling procedure was very laborious and resulted in ∼1 μl lymph after four to five individual cannulations per mouse. Clearly, this technique is not suitable for regular use with animal models or translatable to humans given that the fluid yield was extremely low, and the procedure is invasive and very labor intensive.

Tissue centrifugation and elution are alternative methods when prenodal lymph is not available. A major advantage with the centrifugation method is that native IF is isolated, allowing for direct quantification of mediators. Tissue elution is advantageous in that the method can be applied in situations where the amount of tissue is limited, as shown here for human intestinal mucosa biopsies. A major advantage to both techniques is that individual tissues can be sampled and used for analysis without pooling. The centrifugation technique was originally developed for IF isolation from rat tumors and skin^18^, and although it has subsequently been applied to other organs (for review see^15^), to our knowledge the method has not been evaluated for IF isolation in the gut. We have previously concluded that if the method can be validated for an organ/tissue, the centrifugation method can be considered a reference method for native IF isolation and to determine local production of substances.^15^ Depending on the intestinal segment, these tracer experiments showed that intracellular fluid contributed to ∼10-40% of the isolated fluid (Figure 1D).

The amount of IF that can be collected from each sample is partially related to the size of the initial tissue segment. Therefore, the centrifugation technique has limitations when used with small tissue segments (mouse, human biopsies). Prospectively, however, it should be possible to obtain a larger tissue sample from resected intestinal tissue obtained during surgery, e.g., in patients suffering with IBD or intestinal malignancies. For small biopsies from colonoscopies, we have also explored the use of tissue elution for isolation of intestinal IF. This versatile technique was originally developed to isolate potential biomarker proteins from the interstitium of tumors^29^. As discussed previously^23^, the tissue preparation by sectioning described in some elution methods may result in the addition of intracellular fluid to the eluate, thereby making it difficult to differentiate between interstitial and cellular origin of the assayed substances, a potential effect that was counteracted here by not sectioning the individual intestinal samples but leaving them otherwise intact in the buffer solution. The fact that the sample is added to a buffer during the elution procedure may pose a particular challenge when the exact concentration is required to determine whether a substance is produced locally, i.e., is an inherent part of the microenvironment, or is delivered to the tissue via the circulation.^23^ However, this limitation can be overcome by back-calculation after careful consideration of dilution factors. Nevertheless, we have been able to show consistent results from elution and centrifugation experiments performed in rats and mice. Importantly, however, the proteomic database used for analysis was focused on host-specific proteins. Analysis of microbial proteins may have told a different story. Taken together, the centrifugation and elution methods have their inherent limitations but complement each other in providing access to the intestinal interstitial fluid phase and microenvironment.

Studies have shown that the interactions between immune cells and the microbiome within the GI epithelium, directly or mediated by microbial byproducts, have broader implications for the host immune system in health and disease.^30^ While the composition of immune cells in the gut^22^, in the draining lymph nodes^31^, and in distal organ systems^32^ has been the subject of several investigations, the content of the intestinal mucosa IF, with the notable exception discussed above^28^, has been largely ignored. Interactions between humans and their intestinal inhabitants often focus on correlating phenotypic changes in the host with microbial or metabolomic signatures in feces, which often contain a variety of undigested food residues.^33^ Others have also recognized the need for alternative methods to re-evaluate the microbiome in the context of its site-specific interactions with the host. Indeed, Zmora *et al.* demonstrated that the microbiome of fecal pellets is only partially representative of the contents captured by site-specific sampling of the GI tract.^34^ In a proof-of-principle study, De la Paz *et al*. demonstrated that an ingestible biosensing system can be used for *in situ* monitoring of the GI tract in a porcine model, with the goal of monitoring site-specific host-microbiome interactions in future.^35^ Alternatively, the capture of IF within the GI tract may aid in our understanding of how uptake of certain molecules into the host tissue microenvironment can impact health and disease.

### Limitation

Above, we have discussed the limitations of the centrifugation and elution methods per se. In addition, our study was intended to be a mythological proof-of-principle and feasibility study, not a clinical trial. Therefore, we analyzed only three IBD patients and four healthy controls as use cases. For a more comprehensive understanding of the mucosal microenvironment, larger clinical studies are needed, which we have recently initiated (*InFlame* study; DRKS00031203). The small size of a biopsy of only about 9 mg made it impossible to obtain native fluids with the centrifugation method and forced us to use the elution method, which allows only indirect quantification. The procedure for obtaining human intestinal biopsies also has some limitations. Routinely, a bowel lavage is performed prior to the clinical colonoscopy. Thus, absolute levels of certain microbial-derived metabolites such as SCFA, that are dependent on a dietary substrate, may be affected by a potential lack of continuous substrate supply due to the bowel cleansing. However, we expect that the confounding procedure would be systematic in all study participants.

### Translational perspective

More than 2,000 years ago, Hippocrates claimed that ‘all disease begins in the gut’. The ability to collect IF from tiny human intestinal biopsies and directly measure the microenvironment at the site-of-action, where microbes and mucosal immune cells reside, will overcome the limitations of indirect stool sample analysis. Our pilot proof-of-principle cytokine data pinpoint the individualized inflammatory signatures within the tissues of IBD patients. Large studies such as the *CANTOS* trial^36^ in patients at risk for CVD with thousands of patients with high hsCRP have shown that anti-IL-1β treatment improves cardiovascular outcomes. Several aspects of the *CANTOS* trial inform our current work. The hsCRP-IL-1b-inflammasome axis was critical for a successful cardiovascular outcome, although large numbers of patients were needed to demonstrate benefit. Anti-inflammatory treatment was effective only in a subset of patients at risk for CVD, although identification of specific risk indicators may help to reduce treatment failures. To eventually provide patients with personalized medicine, we need to be able to identify which patients would benefit from targeted interventions. Given that our new approach can be used to identify patients with unique cytokine profiles, targeted recruitment could improve the efficacy of interventions and reduce the number needed to treat. Since the diversity of presentations and drivers of IBD flares presents a challenge, we suspect that the IF isolation from the GI tract may help us to overcome, supporting the concept of personalized medicine.

## AUTHOR CONTRIBUTIONS

EGA, LMH, DNM, and HW contributed to study design. EGA, LMH, SMK, ON, TVK, AY, HB, TUPB, MIW, and OT contributed to data collection. GZ, LMH, AK and BS collected the biopsies and were responsible for the clinical procedures. EGA, LMH, DSV, MK, OP, SG, and TVK contributed to data analysis. AW and GG performed the multiplex cytokine analyses. VMcP, RK, NH, SMK, PM, BM, NW, SKF, HW, and DNM provided expert opinion and supervision. EGA, LMH, DNM, and HW contributed to manuscript preparation.TRR241 IBDome Consortium: Imke Atreya^1^, Raja Atreya^1^, Petra Bacher^2,3^, Christoph Becker^1^, Christian Bojarski^4^, Nathalie Britzen-Laurent^1^, Caroline Bosch-Voskens^1^, Hyun-Dong Chang^5^, Andreas Diefenbach^6^, Claudia Günther^1^, Ahmed N. Hegazy^3^, Kai Hildner^1^, Christoph S. N. Klose^6^, Kristina Koop^1^, Susanne Krug^3^, Anja A. Kühl^3^, Moritz Leppkes^1^, Rocío López-Posadas^1^, Leif S.-H. Ludwig^7^, Clemens Neufert^1^, Markus Neurath^1^, Jay Patankar^1^, Magdalena Prüß^3^, Andreas Radbruch^4^, Chiara Romagnani^3^, Francesca Ronchi^6^, Ashley Sanders^3,8^, Alexander Scheffold^2^, Jörg-Dieter Schulzke^3^, Michael Schumann^3^, Sebastian Schürmann^1^, Britta Siegmund^3^, Michael Stürzl^1^, Zlatko Trajanoski^9^, Antigoni Triantafyllopoulou^5,10^, Maximilian Waldner^1^, Stefan Wirtz^1^, Sebastian Zundler^1^

^1^Department of Medicine 1, Friedrich-Alexander University, Erlangen, Germany

^2^Institute of Clinical Molecular Biology, Christian-Albrecht University of Kiel, Kiel, Germany.

^3^Institute of Immunology, Christian-Albrecht University of Kiel and UKSH Schleswig-Holstein, Kiel, Germany.

^4^Charité – Universitätsmedizin Berlin, corporate member of Freie Universität Berlin and Humboldt-Universität zu Berlin, Department of Gastroenterology, Infectious Diseases and Rheumatology, Berlin, Germany

^5^Deutsches Rheuma-Forschungszentrum, ein Institut der Leibniz-Gemeinschaft, Berlin, Germany

^6^Charité – Universitätsmedizin Berlin, corporate member of Freie Universität Berlin and Humboldt-Universität zu Berlin,Institute of Microbiology, Infectious Diseases and Immunology

^7^Berlin Institute für Gesundheitsforschung, Medizinische System Biologie, Charité – Universitätsmedizin Berlin

^8^Max Delbrück Center für Molekulare Medizin, Charité – Universitätsmedizin Berlin

^9^Biocenter, Institute of Bioinformatics, Medical University of Innsbruck, Innsbruck, Austria.

^10^Charité – Universitätsmedizin Berlin, corporate member of Freie Universität Berlin and Humboldt-Universität zu Berlin, Department of Rheumatology and Clinical Immunology, Berlin, Germany

#EGA and LMH contributed equally

†DNM and HW contributed equally

## ACKNOWLEDGEMENT

We thank J. Czychi, I. Kamer and Dr. Trude Skogstrand for their technical assistance.

## SUPPLEMENTARY METHODS

### Animal protocols

Experiments were carried out in both Berlin, Germany at the ECRC as well as the Bergen, Norway at the University of Bergen. Experiments carried out at the ECRC were approved by the local ethics committee and were compliant with the German and European legal animal protection standards (C57BL/6J mice: X9009/18, SD rats: Y9004/18). Mice and rats were provided unlimited access to food and water, and maintained on a 12:12, light: dark cycle throughout the experiment. SD rats were aged 19-22 weeks and included both males and females. All mouse experiments were carried out using male C57BL/6J mice aged 8-13 weeks. Mice were euthanized by isoflurane anaesthesia. Experiments carried out at the University of Bergen were approved by the AAALAC International Accredited Animal Care and Use Program (approval ID #10508 and #13922) and complied with the regulations put forth by the Norwegian State Commission for Laboratory Animals. Specificities regarding the use of animals for experimentation is described in detail below. Wherever applicable, we followed the *ARRIVE* guidelines.

### Human protocols

Written informed consent was obtained from all patients and healthy volunteers. All experiments were approved by the institutional board of the Charité-Universitätsmedizin Berlin (EA1/200/17) and conducted in accordance with the principles set out in the declaration of Helsinki.

### Centrifugation method

The centrifugation method is outlined in Wiig et al(3). Modifications to the method were made based on expert opinion to create a method for GI tissue centrifugation for use with mouse or rat specimens. To avoid evaporation, these experiments were performed in a humid environment. An out-of-use incubator provided by the University Clinic in Erlangen was used as a humid chamber. The humid chamber was brought to 100% humidity at room air temperature and was lined with a water repellant surface for sample preparation. The schematic in Fig. 1A illustrates the centrifugation technique as it is described here, though the incubator is not shown. Prior to experimentation one 5 mL Safety cap reaction vessel (Ratiolab) per specimen was weighed with a cut piece of nylon weaved mesh netting (pore size ∼15 × 20 μm, Burmeister AS). The size of the netting was adjusted depending on the animal model used to account for the size difference of their respective GI tissues. A 9 x 9 cm piece of netting was used for rat tissues and 7 x 7 cm for mice tissues. The mesh netting suspends the tissue above the apex of the 5 mL tube while the IF descends into the bottom of the tube during centrifugation. To prevent puncturing delicate GI tissue during preparation, small pieces of 0.5 mm ROTILABO ® silicone hose (Carl Roth) were used to cover sharp surgical tools.

All samples of GI tissue were collected promptly after the sacrifice of each mouse or rat. The distal esophagus to the distal colon was removed, placed immediately in a closed petri dish, and placed inside the humid chamber to avoid evaporation. The sectioning approach varied based on the goal of the experiment and the animal model used and is listed alongside each figure as appropriate. In general, the five segments were collected from the colon, cecum, ileum, jejunum, and duodenum in mice. From rats as the GI tract yields significantly more tissue, non-cecal segments were split into multiple segments for practical reasons. Samples from these segments were not pooled but used separately for analyses. Segments were ligated and cut between the ligations, placed onto mesh netting, then gently positioned inside the 5 mL tube with the netting tucked under the tube cap to suspend the tissue above the tube apex. During preparation, pieces of tissue not yet prepared were kept inside a closed petri dish within the humid chamber. Once all tubes were filled, they were centrifuged together for 10 minutes at 400 x g. After centrifugation, GI tissue and mesh was removed and set aside. Tubes containing centrifuged IF were weighed and ddH_2_0 was added 1:10 as diluent. Diluted IF was then aliquoted and frozen at −80°C.

### Elution method

The elution approach is described by Wiig et al(3), and was modified for use with GI tissue as shown in Fig. 3A. The full GI tract was removed from each mouse or rat as described above and promptly placed in the humid chamber. As described above, surgical tools were sheathed in silicon hose to minimize damage to the gut. Approximately 2-4 cm sections from the colon, cecum, ileum, jejunum, and ileum were excised and separated (unless otherwise stated). The inner surface of each GI segment was flushed with isosmotic elution buffer to remove any remaining fecal matter using a 1.7 x 50 mm Vasofix® Safety IV catheter (B. Braun Medical Industries) without a needle, which was attached to a 5 mL syringe (B. Braun Medical Industries). Segments were then gently manipulated to remove the remaining elution buffer, ligated, and positioned in a pre-weighed 15 mL tube. Each segment was weighed, and isosmotic elution buffer was added in a 1:10 ratio to the weight. Segments were then held at 4°C and rocked gently on a rocker for a defined time (2-48 hours). After the elution was complete GI segments were removed, and the elution IF was aliquoted and frozen at −80°C.

**Figure 3.**
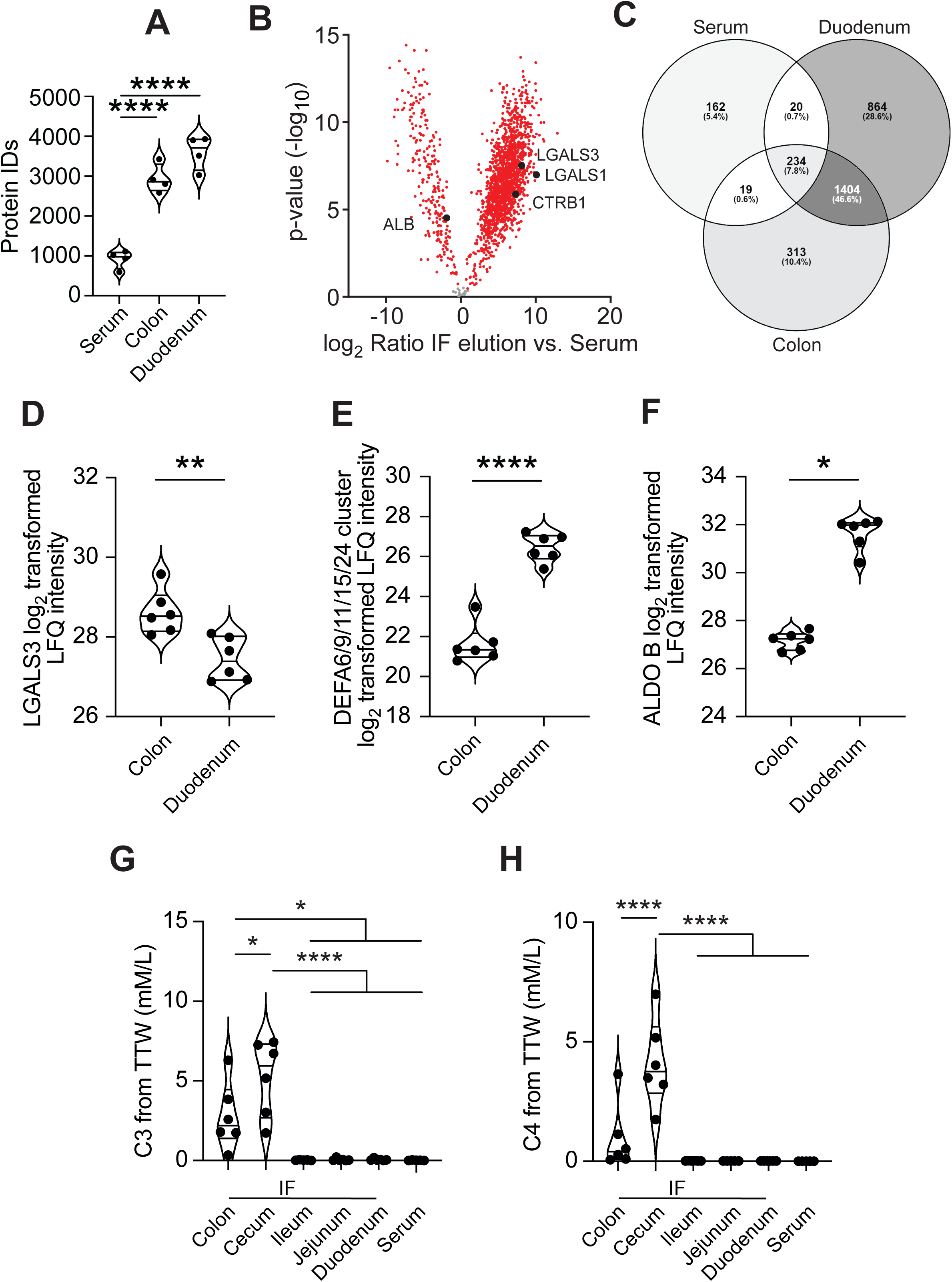
The serum and elution IF proteomes are unique in C57BL/6J mice. Eluate samples (colon and duodenum) from C57BL/6J mice were analyzed by shotgun proteomics and compared to serum samples (n=4 for each condition) for (A-C). Elution IF was collected after 4 hours in a mannitol-based buffer solution. The Mouse Uniprot database 2019 was used for data curation. A) The total number of protein IDs found in individual samples from the colon, duodenum, and serum. B) Proteins which were found to be significantly different from eluate samples compared to the serum. The actual −log_10_ transformed p-values and the log_2_ difference between elution and serum for a given protein are shown as individual dots, and red dots indicate significant p-values after an unpaired two-tailed *t*-test with an FDR correction of 5%. The number of overlapping proteins from serum, duodenal and colonic samples are shown in C). A given protein had to have 100% valid values within at least one group to be included in the comparisons for both (B-C). The Venn diagram was generated using the Venny tool (bioinfogp.cnb.csic.es/tools/venny/index.html). A separate experiment with n=3 C57BL/6J mice and two segments per anatomical segment (n=6 total) was analyzed using shotgun proteomics for (D-E). Elution IF was collected after 2 hours in a mannitol-based solution. D-E) Known proteins of importance in the GI tract were detectable in elution IF. The log_2_ transformed LFQ intensity values for individual proteins LGALS3 (D), the α-defensin (DEFA) cluster 6/9/11/15/24 (E), and ALDOB (F) are shown. For (G-H), samples from C57BL/6J mice (n=6) were eluted for 2 h in a saline-based buffer solution for measurement of SCFA by GC-MS. G) Propionate (C3) and butyrate (H) levels in IF from different segments of the mouse GI tract compared to plasma. Significance was tested using a paired two-tailed t-test for D-E and Wilcoxon for F and an one-way ordinary ANOVA with Tukey’s multiple comparison test for A, G-H. *p≤0.05, **p≤0.01, ***p≤0.001, ****p ≤ 0.0001.

**Figure 4.**
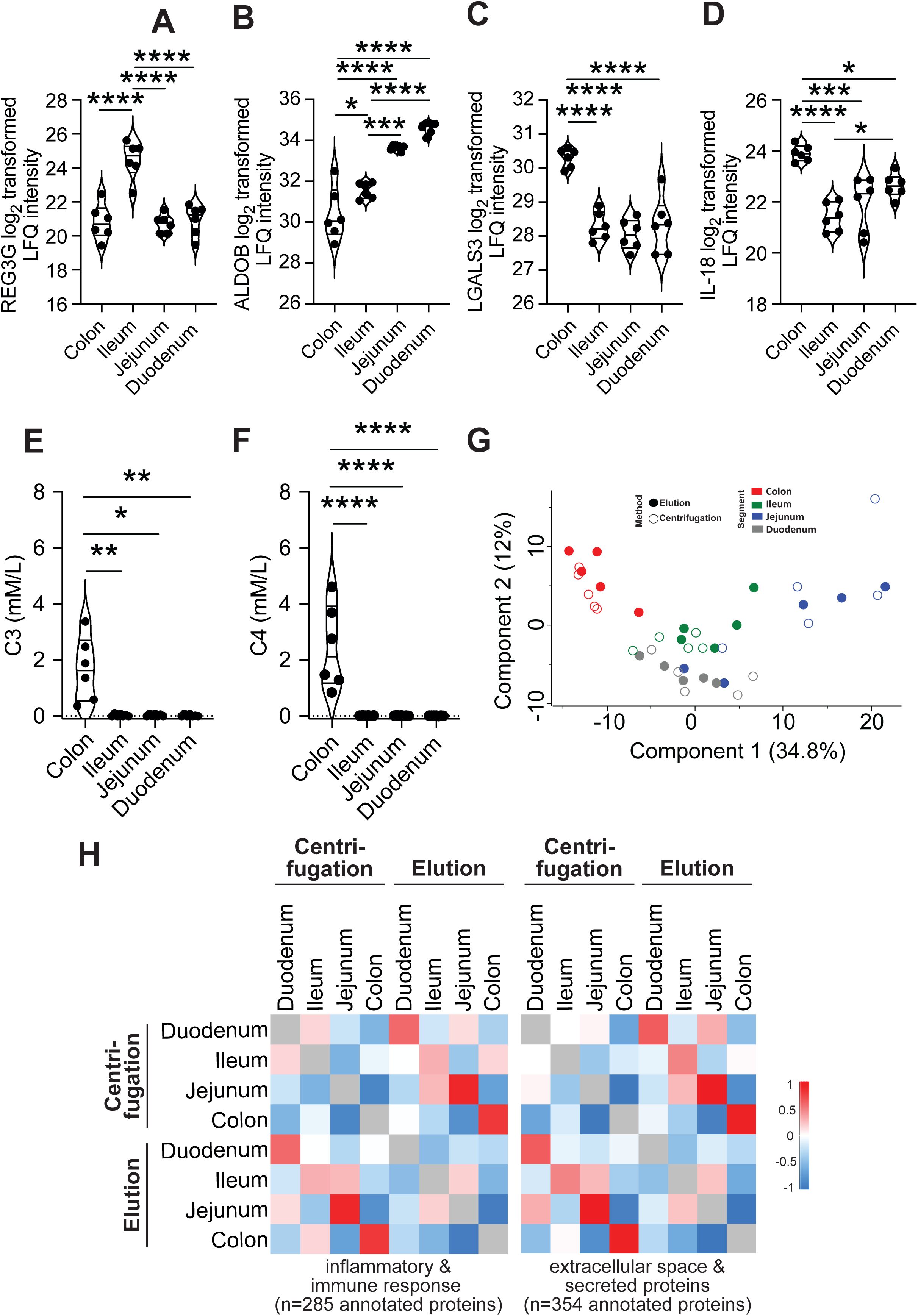
GI-related proteins, cytokines and SCFA expected in IF are similarly enriched in various GI segments in both the elution and centrifugation methods. Centrifugation and elution IF were collected in tandem from SD rats (n=6). Samples were eluted in saline buffer for 2 hours. A-D) Selected candidates (REG3G, ALDOB, LGALS3 and IL-18) were analyzed by mass spectrometry in rat IF isolated by the elution method. E-F) Propionate (C3) and butyrate (C4) levels measured by GC-MS from IF isolated by the elution method. Independent z-scoring was performed within the centrifugation and elution methods, and only proteins which appeared with 100% valid values for at least one segment in both methods were included in the data shown in (G-H, n=5). Selection of inflammatory response and immune response annotations (285 proteins) from the rat Uniprot database is shown by (G) PCA and (H) Pearson’s correlation. Pearson’s correlation of selected extracellular space and secreted proteins (354 proteins) from the rat Uniprot database is shown in (H). In (G), the point fill indicates the method, and the point color indicates the segment of origin (colon, ileum, jejunum, or duodenum). In (H), the mean Pearson’s correlation is shown for segments within and between methods. Significance for A-D was tested by ordinary one-way ANOVA with Tukey’s multiple comparison test. For E-F) Kruskal-Wallis test corrected for multiple comparison by controlling FDR (Benjamini, Krieger, Yekutieli) *p≤0.05, **p≤0.01, ***p≤0.001, ****p<0.0001 (n=5-7).

**Figure 5.**
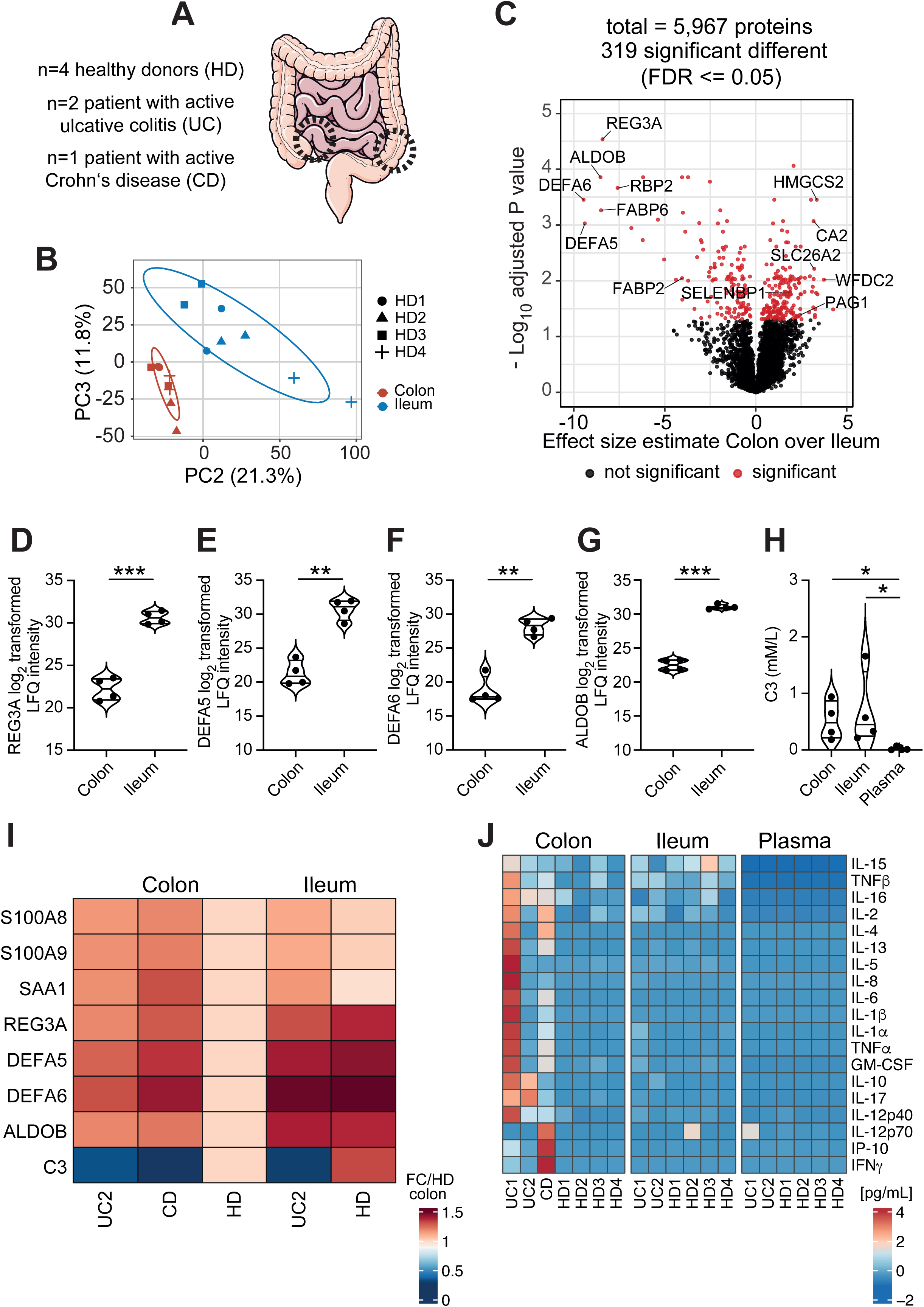
Distinct proteomes and cytokine signature in colon and ileum elution IF samples from healthy human donors and IBD patients. A) The schematic of the human GI tract shows the location of the colon and ileum biopsy sites (dashed circles). The number of healthy donors (HD; n=4), patients with active ulcerative colitis (UC; n=2) or active Crohn’s disease (CD; n=1) included in the study is indicated. Part of the panel A) was drawn with Servier Medical Art by Servier, licensed under a Creative Commons Attribution 3.0 Unported License. B-G, I) Elution IF samples (colon and ileum) from healthy donors and diseased patients were analyzed by label-free proteomics. Elution IF was collected in an isotonic sodium chloride solution for 2 hours. B) Principal component analysis (PCA) of the IF from gut compartments of the colon (red) and ileum (blue) of healthy donors based on 5,967 proteins after data filtering. The second and third PCs represent a cumulative variance of 33.1% and separate the data sets by compartment. Two technical replicates per donor per compartment are shown, and the data were scaled and centered for PCA analysis. C) Differentially regulated proteins between healthy colon and ileum samples. Each dot represents a unique protein ID plotted based on the effect size estimate and adjusted p-value between colon and ileum samples. Red dots represent significant p-values tested with a paired *t*-test and a BH-FDR correction of 5%. Proteins relevant to the GI tract are highlighted. Comparison of selected candidates (REG3A, DEFA5, DEFA6, ALDOB) in colonic and terminal ileal IF of healthy donors (D-G) and IBD patients (I-J) determined by performed label-free proteomics (D-I) or multiplex cytokine assays (J). H) Propionate (C3) levels in colonic, ileal IF of healthy donors compared to plasma. Significance for D-G was tested using a two-tailed Welch’s paired *t*-test corrected for multiple comparisons by controlling for FDR (Benjamini, Krieger, Yekutieli). For H) Kruskal-Wallis test corrected for multiple comparison by controlling FDR (Benjamini, Krieger, Yekutieli) *p≤0.05, **p≤0.001, ***p≤0.0001.

The isosmotic elution buffer used was dependent on intended use of IF and is listed as such, though for the most part isosmotic saline was used. Mannitol elution buffer was required when ion concentrations were to be measured from elution IF. Due to a matrix-effect, saline elution buffer was required for measurement of SCFA from resultant IF. Prepared isotonic 0.9 % sodium chloride solution for injection (B. Braun Medical Industries) was used as saline elution buffer. Mannitol elution buffer was formulated to an osmolarity of between 285-295 mOsm/kg by adding 5 % pure (99.9999% identity) D-mannitol (Sigma-Aldrich) to double-distilled water (18.2 Ωm). Knauer Semi-Micro Osmometer – Type ML was used to test buffers to ensure the correct physiological osmolarity.

For the elution procedure for human biopsies, two biopsies from respective locations were collected during colonoscopy and immediately placed into a pre-weighed tube containing ice-cold isotonic 0.9 % sodium chloride solution. Samples were incubated on a rocking device at 4 °C for 2 hours. Prior to incubation, biopsy weights were determined. After incubation, biopsies were retrieved and transferred into formalin and the interstitial fluid solution was aliquoted and frozen in liquid nitrogen and stored at −80 °C for further analysis.

### Ion Chromatography

Sodium and potassium in the eluted tissue solutions and serum were baseline separated in a 10 minutes 7.5-60 mmol/LMSA gradient at a flow rate of 0.2 mL/min by a Dionex IonPac CS 16-4µm RFIC analytical column (2 × 250 mm, P/N 088582) and guard (2×50 mm, P/N 088583) using a Dionex Integrion HPIC System equipped with a CDRS 600 (2 mm) Cation Electrolytic Suppressor and a high pressure EGC 500 methane sulfonic acid eluent generator cartridge. Thereafter ion content was related to tissue wet weight, dry weight, and water content.

### 51Cr-EDTA tracer experiments

#### Centrifugation

Experiments were performed at the University of Bergen (approval ID #10508 and #13922). The radiolabeled isotope ^51^Cr-EDTA was used as an extracellular tracer. The relative level of the tracer in IF compared with serum was used to assess whether obtained fluid has been diluted by intracellular contents or other compartments. 2 % isoflurane in 100 % O_2_ was used for anesthesia. During the experiment a servo-controlled heating pad at 37°C was used to maintain body temperature. Male C57BL/6 mice (n=7) were anesthetized and both kidney pedicles ligated to prevent tracer escape during the experiment before ^51^Cr-EDTA (∼ 6 million counts in 100 µL isotonic saline) was injected into the tail vein using an insulin syringe. Mice were kept under continuous anesthesia as ^51^Cr-EDTA equilibrated within the extracellular fluid phase for 1 hour. After the equilibration period a blood sample was obtained by cardiac puncture and mice were sacrificed by cervical dislocation. Mice were immediately transferred to a humidity chamber with 100 % humidity at room temperature for the rest of the harvesting procedure. The centrifugation protocol was then performed as described above. A small piece of skin was also removed and centrifuged as described previously(6). ^51^Cr-EDTA from isolated GI IF, skin IF, and serum samples were counted in a gamma counter 1h after collection. Retrieval of ^51^Cr-EDTA in IF was calculated and expressed as the IF/serum count ratio for each sample from each mouse, respectively. This procedure was performed with minimal modifications in SD rats (n=8) in accordance with centrifugation methods section above. Due to larger body size, rats received a higher dose of chromium (∼15 million counts in 100 µL isotonic saline) and the chromium was left to equilibrate for 2 hours (compared to 1hr in mice) prior to sacrifice.

#### Elution

Experiments were performed at the University of Bergen (approval ID #10508 and #13922). ^51^Cr-EDTA was used to assess the rapidity and repeatability inherent to the elution method as it was applied for use with GI tissue in mice. Mice were continuously anesthetized with 2 % isoflurane in 100 % O_2_ for the duration of experimentation until sacrifice and body temperature was maintained at 37°C with the aid of a servo-controlled heating pad. After ligation of both kidney pedicles ^51^Cr-EDTA (∼ 14 million counts in 100 µL isotonic saline) was injected into the tail vein with an insulin syringe of male C57BL/6J mice (n=6). The ^51^Cr-EDTA was equilibrated in the extracellular fluid phase for 1 hour, after which, a blood sample was obtained by cardiac puncture. Mice were then sacrificed by neck dislocation and transferred to a humidity chamber as described above for the rest of the procedure. Elution IF samples were collected as described in the method above using a mannitol-based elution buffer. The samples were placed in a cold room (4°C) on a rocker and after 2, 4, 6, 24 and 48 hours 100 µL from each sample was removed for gamma counting. After 48 hours, all GI segments were removed from the elution buffer and counted. The fraction of ^51^Cr-EDTA eluted was determined by dividing the count within each elution IF sample at each time point with total counts in each corresponding gut segment prior to elution. Extracellular fluid volume was determined for each gut segment as the plasma equivalent space of ^51^Cr-EDTA.

### Total tissue water determination

Total tissue water (ECV and ICV) of a given GI segment was characterized from C57BL/6J mice (n=7) by removing the GI tract as described above. GI tissue was segmented into segments of colon, cecum, ileum, jejunum, and duodenum that were then cut lengthwise to remove fecal contents. Segments were lightly rinsed with 1x DPBS (Gibco), and lightly blotted to remove excess fluid. Cleaned segments were weighed and dried in ceramic dishes in the UF450 drying oven (Memmert GmbH) to dry at 70°C. Tissue segments were weighed after 36 and 100 hours to ensure that all fluid had left the tissue. Weights did not change beyond 36 hours, as each tissue segment had been dried completely. The GI segment dry weight was divided by the respective wet weight to yield the fractional contribution of fluid compartments to the overall tissue weight.

### Sample preparation and protein digestion for proteomics

#### Mouse and rat IF samples

A quantity of 20 µg of mouse or rat IF was lysed in a 2x SDC-buffer containing 2% Sodium Deoxycholate (Sigma-Aldrich), 20 mM Dithiothreitol (Sigma-Aldrich), 80 mM Chloroacetamide (Sigma-Aldrich), and 200 mM Tris-HCl (pH 8). The lysates were heated for 10 minutes at 95°C and then subjected to digestion with endopeptidase LysC (Wako) and sequence grade trypsin (Promega) at a protein:enzyme ratio of 50:1. Digestion took place overnight at 37°C.

#### Human tissue samples

For human IF samples, the same 20 µg amount was lysed in the 2x SDC-buffer as described. Protein digestion was carried out using trypsin (Promega) at an enzyme-to-protein ratio of 1:20, with the digestion conducted overnight at 37°C.

### Proteomics LC-MS

#### Rat and mouse samples

The resulting peptides were desalted using stage-tips (24) and then subjected to reversed phase liquid chromatography coupled with mass spectrometry (LC-MS). About 2 µg of peptides were injected into an EASY-nLC 1200 system (Thermo Fisher Scientific) for separation. For rat experiments, a Q Exactive HF-X orbitrap mass spectrometer (HFX, Thermo Fisher Scientific) was utilized, while a Q Exactive Plus (QE+, Thermo Fisher Scientific) mass spectrometer was used for mouse experiments.

Both mass spectrometers were operated in a data-dependent acquisition (DDA) mode. Full scans were conducted at 60K resolution (HFX) or 70K resolution (QE+), followed by data dependent MS2 scans of the top 20 precursors. The MS2 scans were performed at 15K resolution (HFX) or 17.5K resolution (QE+), with ion count targets of 1e5 or 5e4 and isolation windows of 1.3 m/z or 1.6 m/z for HFX or QE+, respectively. A dynamic exclusion of 30 seconds was applied for both setups.

#### Human samples

For human samples, peptides were desalted and separated using the EASY-nLC 1200 system, followed by analysis on an Exploris 480 orbitrap mass spectrometer (Thermo Fisher Scientific) operating in data-independent acquisition (DIA) mode. Full scans were conducted at 120K resolution, followed by MS2 scans with variable window widths. Stepped normalized collision energy settings (26, 29, 32) were used, and the MS2 resolution was set to 30K.

### Proteomics data analysis

#### Rat and mouse samples

Raw data were processed using the MaxQuant software package (v1.6.10.43 for mouse samples, v1.6.3.4 for rat samples). (25) MS2 spectra were searched against a mouse or rat decoy UniProt database (MOUSE.2019-07; RAT.2019-07) using the Andromeda search engine. (26) Variable modifications included oxidation (M), N-terminal acetylation, and deamidation (N and Q), while carbamidomethylated cysteine was considered a fixed modification. Peptide length was restricted to a minimum of 7 amino acids, with a maximum of three missed cleavages allowed. The false discovery rate (FDR) was set to 1% for peptide and protein identifications. The integrated label-free quantification algorithm was activated. (27) Unique and razor peptides were considered for quantification and the match-between-runs algorithm was turned on. For further data analysis the Perseus software package (v1.6.2.1) was consulted. (28) MaxLFQ intensity values were used for quantification. (27) Reverse hits, contaminants and proteins only identified by site were filtered out. A minimum of three valid values was required in at least one experimental group. Where appropriate, missing values were imputed by random draw from Gaussian distribution with 0.3*SD and downshift of 1.8*SD of the observed values per sample. Euclidean distances were used for all cluster-based analyses along an axis. Pearson’s correlation was used to assess the similarities between given samples either individually or by the group mean. Principal component analysis (PCA) with Euclidean distances was used to assess similarities between samples. An FDR correction of 5% was applied for statistical testing unless otherwise stated.

#### Human samples

Raw data for human samples were analyzed using DIA-NN version 1.8.1 (29) in library-free mode. The analysis involved searching against a human Uniprot fasta file with isoforms (HUMAN.2022-03). Subsequent data analysis was carried out in R using the report.tsv output. Precursors were filtered based on Q-values (Q.value ≤ 0.01, Protein.Q.value ≤ 0.01, Lib.Q.Value ≤ 0.01), and LFQ intensities were averaged across precursors and collapsed to protein group level. Protein groups were filtered for a minimum of 75% valid values across all conditions before applying downshift imputation. Differential expression analysis employed the paired Welsh test, and the Benjamini-Hochberg correction was used for multiple hypothesis testing.

### Measurement of SCFA using GC-MS

SCFA from serum, and IF samples were measured as per the method described previously with minor modifications.(30) Briefly, 90 µL of murine serum or IF were extracted by shaking samples in 100 µL diethyl ether and 10 µL HCl at 25°C for 30 min. To account for low sample volume, predilution of samples with diluent of ddH_2_O was used where appropriate. Samples were then centrifuged for 5 min at 1500 x g, after which time, 50 µL of the ether phase was transferred into GC-MS glass vials. 10 µL MTBSTFA was added for derivatization and samples were held at 80°C for 30 min, and subsequently held overnight incubation at room temperature. Human serum samples were handled equally. For human IF samples, 45 µL of sample were extracted with 45 µL diethyl ether, 5 µL HCl and 25 µL of 60% (w/v) NaHSO_4_. Derivatization was performed on 20 µL ether phase with 4 µL MTBSTFA. Spike in of 100 µM crotonic acid was used as an internal control for all samples. 1000 µM sodium acetate was added to murine serum samples to increase the signal above background, hence 1000 µM was subtracted after quantitation from these samples. A dilution series of Volatile Free Acid Mix (CRM46975, Sigma Aldrich) and pure ddH_2_O were prepared in parallel and measured with each run for absolute quantification. SCFA calibration curve in a saline-based elution buffer was similar to in ddH_2_0, therefore this buffer was used for all experiments. GC-MS analysis was performed on a Thermo Scientific™ Q Exactive™ hybrid quadrupole Orbitrap mass spectrometer, coupled to a Thermo Scientific™ TRACE 1300 Series gas chromatograph and a Thermo Scientific™ TriPlus RSH Autosampler. For murine samples, 1 µL and for human samples, 4 µL were injected, each with a split of 1:10. Gas-chromatographic separation was performed with an initial temperature of 68°C, held for 2 minutes, followed by a 7 °C/minute ramp until 150 °C and a final 50 °C/minute ramp until 300 °C, held for 2 minutes. Full MS was acquired at a resolution of 60,000 and a scan range of 65 to 600 m/z. Thermo Scientific™ Xcalibur™ Quan Browser Software was used for data analysis. If extraction was not successful or internal standards were not measurable these samples were removed from the analysis.

### Multiplex cytokine analysis of IF

To characterize the intestinal cytokine profile, IF isolated from human biopsies as described above were analyzed using the human V-Plex Plus Human Cytokine kit (IL-1α, IL-1β, IL-2, IL-4, IL-5, IL-6, IL-8, IL-10, IL-12p40, IL-12p70, IL-13, IL-15, IL-16, IL-17, TNFα, TNFβ, IP-10, INFγ) from Meso Scale Diagnostics according the manufacturer’s instructions. Analysis was performed at the Immunological Study Lab of CheckImmune GmbH on a Mesoscale Discovery platform. For each sample, the respective value of the electrochemiluminescence (RFU-Signal) of the analyte concentration was calculated based on a calibration curve. Measurements of the Meso Scale Diagnostics assay were performed and evaluated in accordance with the ICH-GCP Guideline ‘Validation of analytical procedures’.

### Cytokine and SCFA calculations for tissue fluid space of biopsies

To determine the expected concentration of respective cytokine or SCFA within the total tissue fluid space of eluted biopsies, the following calculations were performed. First, the concentration of cytokine or propionate per gram of tissue (μM/g or pg/g) was calculated using the following formula:

*(Concentration (μM or pg/mL) determined in the elution IF x total elution fluid volume (mL)) / tissue weight (g)*.

Next, the concentration of cytokine or propionate in the total tissue water (μM/mL or pg/mL) was calculated using the following formula:

*Concentration of cytokine or propionate in the tissue (μM/g or pg/g) / 0.79 (mL/g)*.

### Statistical analysis

Statistical methods applied were dependent on the experimental design and are listed appropriately alongside each figure. All analyses were performed in a blinded manner. Data were examined for outliers using the ROUT outlier test. Outliers were removed when appropriate. Where when sequential measurements were taken from one biological sample, repeated-measures one-way ANOVA with Tukey’s multiple comparisons test was used given that these data are considered paired. To assess differences between segments or measurements that were normally distributed and not paired, ordinary one-way ANOVA with Tukey’s multiple comparisons test was used. When the data were not normally distributed the Kruskal-Wallis test corrected for multiple comparison by controlling FDR (Benjamini, Krieger, Yekutieli) was used. Ordinary one-way ANOVA with Dunnett’s post-hoc test was applied when comparing multiple groups to a reference value. Unpaired two-tailed Student’s t-test and two-way ANOVA testing were used depending on the variables being interrogated. P values of less than or equal to 0.05 were considered statistically significant. Statistics particular to the proteomics methods are listed within the respective methods section above. All other statistical analyses were performed using GraphPad Prism 9. The analysis plan was developed before the start of the study. All study data are available on request.

## SUPPLEMENTARY LEGENDS

**Figure S1.**
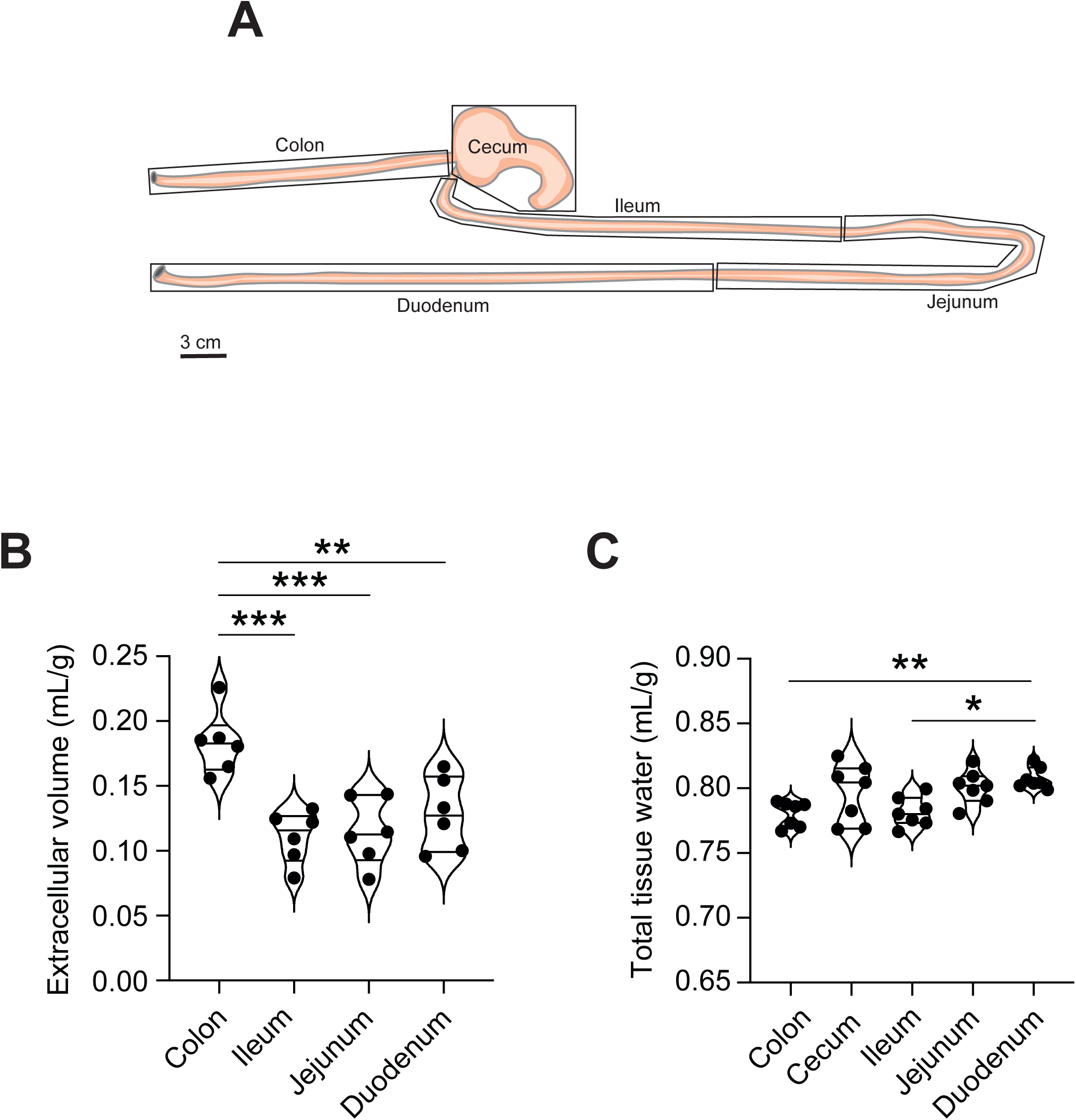
Elution-based extracellular fluid isolation within the GI tract of C57BL6/J mice. A) Schematic of the tissue segments harvested for analysis from C57BL/6J mice (the cecum was used only in (C)). B) Extracellular fluid volume was determined by counting ^51^Cr-EDTA for each intestinal segment compared to the plasma equilibration of the tracer (n=6). In (C) the total tissue water is shown from C57BL/6J mice (n=7). Significance was tested using an ordinary one-way ANOVA with Tukey’s multiple comparison test. *p≤0.05, **p≤0.01, ***p≤0.001.

**Figure S2.**
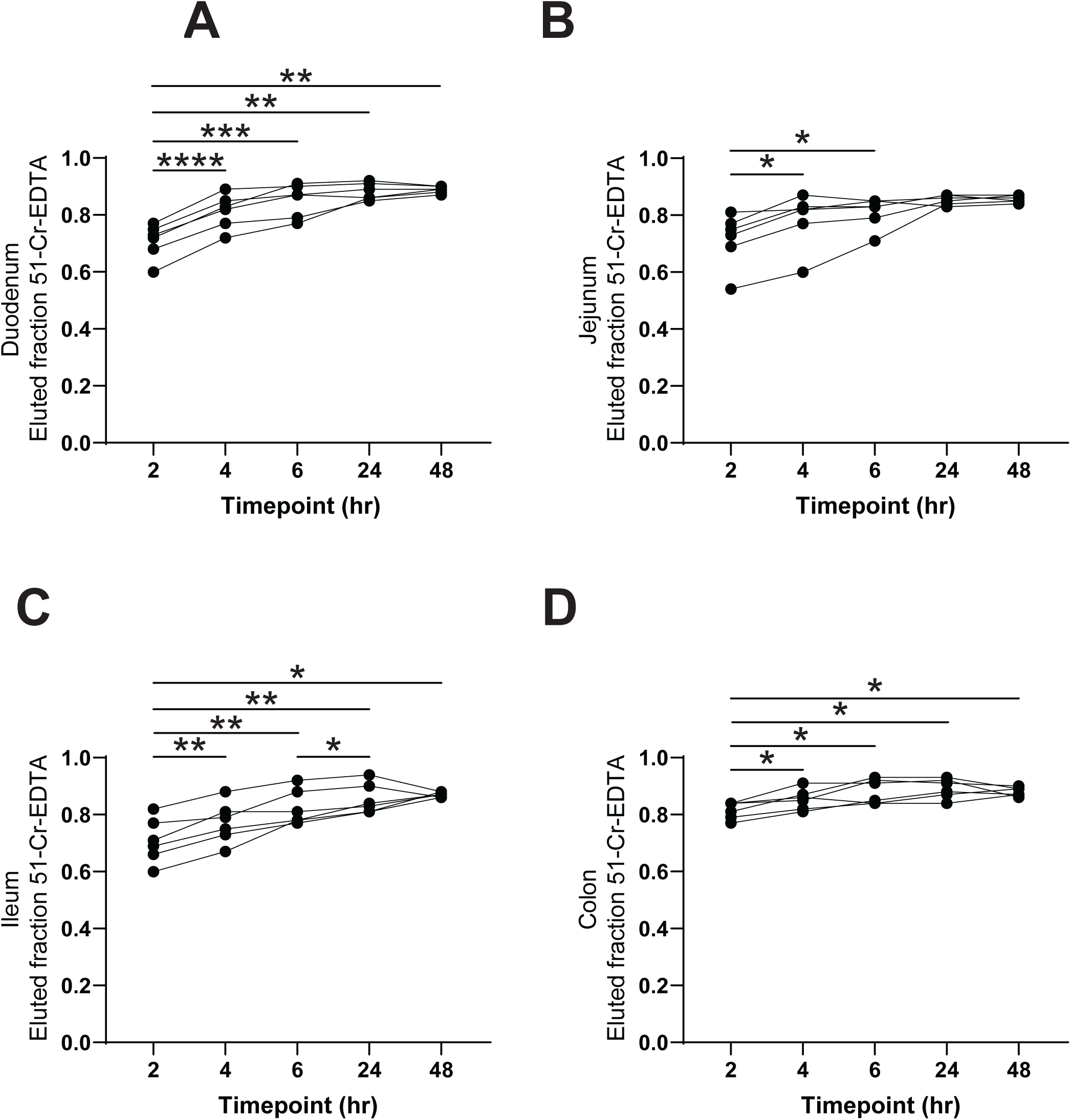
^51^Cr-EDTA elution indicates the rapid equilibration of this extracellular tracer with the surrounding elution buffer. A) The extracellular tracer ^51^Cr-EDTA was used to determine the rate at which the elution fluid equilibrates with the IF from C57BL/6J tissue. In (A-D), at each time point, 100 µL of sample was taken from the eluted solution for gamma counting. The eluted fraction for each segment was determined by dividing the counts in the eluted sample at each time point by the total counts in the corresponding intestinal sample prior to elution. Data (n=6 for all conditions) were tested using repeated measures one-way ANOVA with Tukey’s multiple comparisons test and significant post hoc comparisons are shown. *p≤0.05, **p≤0.01, ***p≤0.001, ****p≤0.0001.

